# An interplay between reaction-diffusion and cell-matrix adhesion regulates multiscale invasion in early breast carcinomatosis

**DOI:** 10.1101/566612

**Authors:** Dharma Pally, Durjay Pramanik, Ramray Bhat

**Author notes:** Equal contribution (Dharma Pally, Durjay Pramanik).

## Abstract

The progression of cancer in the breast involves multiple reciprocal interactions between malignantly transformed epithelia, surrounding untransformed but affected stromal cells, and the extracellular matrix (ECM) that is remodelled during the process. A quantitative understanding of the relative contribution of such interactions to phenotypes associated with cancer cells can be arrived at through the construction of increasingly complex experimental and computational models. Herein, we introduce a multiscale 3D organo-and patho-typic model that approximates, to an unprecedented extent, the histopathological complexity of a tumor disseminating into its surrounding stromal milieu via both bulk and solitary motility dynamics. End-point and time-lapse microscopic observations of this model allow us to study the earliest steps of cancer invasion as well as the dynamical interactions between the epithelial and stromal compartments. We then construct an agent-based Cellular Potts model that incorporates constituents of the experimental model, as well as places them in similar spatial arrangements. The computational model, which comprises adhesion between cancer cells and the matrices, cell proliferation and apoptosis, and matrix remodeling through reaction-diffusion-based morphogen dynamics, is first trained to phenocopy controls run with the experimental model, wherein one or the other matrices have been removed. The trained computational model successfully predicts phenotypes of the experimental counterparts that are subjected to pharmacological treatments (inhibition of N-linked glycosylation and matrix metalloproteinase activity) and scaffold modulation (alteration of collagen density). Our results suggest that specific permissive regimes of cell-cell and cell-matrix adhesions operating in the context of a reaction-diffusion-regulated ECM dynamics, promote multiscale invasion of breast cancer cells and determine the extent to which they migrate through their surrounding stroma.

## Introduction

Within physiologically functioning tissues and organs, cells constantly interact with their surrounding extracellular matrix (ECM). This interaction is complex and is essential for organ development and homeostasis (Bhat and Bissell, 2014; Bhat and Pally, 2017). Aberrant alterations that affect cell-ECM interactions aid in the progression of pathologies like cancer (Nelson and Bissell, 2005; Simi et al., 2018). In normal mammary glands and breasts, luminal epithelial cells are surrounded by a layer of myoepithelial cells that secrete basement membrane (BM): a sheet-like ECM rich in laminin and non-fibrillar collagens. Such mammary epithelial architectures are surrounded by stromal ECM that is rich in fibrillar matrix proteins such as collagens and connective tissue cells such as fibroblasts, macrophages, adipocytes. In breast cancer, this architecture is lost: the lumen is filled with proliferating apolar transformed epithelia, myoepithelia are absent and the BM is ultimately breached by invading cells (Polyak and Kalluri, 2010). The stroma shows degradation of ECM, fibrosis, leucocytic infiltration, neo-angiogenesis and lymphangiogenesis (Dumont et al., 2013; Orimo et al., 2005; Wiseman and Werb, 2002). Malignant transformation results in a recalibration of existent interactions and the origination of novel ones between constituents of the tumor microenvironment. Studying and quantifying the contribution of a given interaction to the progression phenotype of cancer spatiotemporally is a challenge, as our histo- and bio-chemical analyses are limited to distinct stages of breast cancer from various patients. Three-dimensional (3D) organotypic and pathotypic cultures of cancer cell lines and primary patient cells have helped extend our understanding of the molecular mechanisms underlying cancer (Torras et al., 2018; Weinhart et al., 2019). 3D cultures are approximations of the histopathological complexity of *in vivo* tumor microenvironments; most current models involve embedding cancer epithelia within a natural or tunable synthetic matrix scaffolds(Balachander et al., 2015; Bhat et al., 2016; Furuta et al., 2018) (more complicated versions comprise efforts to mimic the perivascular and endothelial metastatic niches (Carlson et al., 2019; Ghajar et al., 2013) as well as efforts to engineer platforms consisting of multiple organs-on-a-chip reviewed by (Zhao et al., 2019)).

Non-cancerous and malignant breast cell lines, when cultured in reconstituted basement membrane matrix (rBM), cluster into discrete morphologies that have been described as ‘round’, ‘mass’, ‘grape’ and ‘stellate’ in increasing order of aggressiveness (Kenny et al., 2007). The round phenotype is characteristic of untransformed cells that form growth-arrested acinar-like multicellular clusters with basoapical polarity, and a lumen. Mass and grape phenotypes are characteristic of malignantly transformed epithelia which mimic carcinoma-in-situ or indolently progressive cancers with cells that have completely lost their polarity. The stellate phenotype is typical of highly metastatic cancer cells that actively migrate, although in a collective manner, into and through rBM matrices. Using such 3D assays of cells embedded in rBM, it is possible to study the role of specific expressed proteins in regulating the adhesion between cancer cells, or with ECM proteins such as laminins. However, such culture frameworks are inadequate for investigations into the dynamics of spatial transition of cells between two matrix microenvironments that have distinct rheological properties, such as the non-fibrillar BM-like microenvironment and its fibrillar collagen-like types (Fig S1A and B shows scanning electron micrographs of non-fibrillar rBM and fibrillar Type 1 collagen matrices). In addition, it is unfeasible to design experiments to observe the phenotypic consequences of an exhaustive exploration of interaction space within a multicomponent biological system.

The second limitation can especially be mitigated by adopting a computational approach and simulating the progression of cancer-like phenotypes for a diverse range of interactive parameter combinations. Agent-based models, particularly the Cellular Potts model (CPM) have been shown to be useful for such efforts (Swat et al., 2012; Zhang et al., 2011). For example, the deployment of proteolytic and non-proteolytic mode of cancer migration through collagenous scaffolds, or between solitary and collective cell invasion has been well-elucidated using *in silico* approaches (Kumar et al., 2016). However, to the best of our knowledge, no theoretical model has explicitly explored the transitioning dynamics, and the consequences thereof, of cancer cells moving between dissimilar matrix-microenvironments. Moreover, while the dynamical role of individual mesoscale physicochemical processes have been well studied in cancer and development (Grant et al., 2004; Pantziarka, 2016; Zhang et al., 2011), whether their combined deployment constrains or widens the phenotypic reaction norm: the spectrum of discrete phenotypes achievable by cancer cells, has not been investigated using such a computational effort.

In this paper, we present a unified experimental-computational framework to investigate the interactions between cancer epithelia and spatially compartmentalized extracellular matrix microenvironments. The experimental model allows us to break down the phenomenon of cancer cell migration into cellular interactions with the BM, their remodelling of the same, their transition from BM to Type 1 collagen, and the subsequent remodelling of, and migration within, Type 1 collagen. We closely train the computational agent-based model on experimental results. The computational model successfully predicts results of the cancer epithelia upon pharmacological perturbations or scaffold modification. The trained theoretical model also predicts that emergent interplay between reaction-diffusion and cell-matrix adhesion can explain the diversity in the extent and mode of invasion of breast cancer cells.

## Materials and Methods

### Cell culture

MDA-MB-231 cells were maintained in DMEM:F12 (1:1) (HiMedia, AT140) supplemented with 10% fetal bovine serum (Gibco, 10270). MCF-7 cells were grown in DMEM (HiMedia, AT007F) supplemented with 10% fetal bovine serum. HMLE cells were a kind gift from Dr Robert Weinberg, Harvard Medical School and Dr Annapoorni Rangarajan, Indian Institute of Science and were cultured in DMEM:F12 (1:1) supplemented with 1% fetal bovine serum, 0.5 μg/mL Hydrocortisone (Sigma, H0888), 10 μg/mL Insulin (Sigma, I6634) and 10 ng/mL human recombinant epidermal growth factor (HiMedia, TC228). All the cells were cultured in a 37°C humidified incubator with 5% carbon dioxide.

### Preparation of cancer cell clusters

Normal/cancer cells were trypsinized using 1:5 diluted 0.25% Trypsin & 0.02% EDTA (HiMedia, TCL007). 30,000 cells per 200 μL of defined medium (Blaschke et al., 1994) supplemented with 4% rBM (Corning, 354230) were cultured on 3% polyHEMA (Sigma, P3932) coated 96 well plate for 48 hours in a 37°C humidified incubator with 5% carbon dioxide.

### 3D invasion assay

rBM-coated clusters were collected into 1.5 mL tubes, centrifuged briefly and supernatant is removed. Acid-extracted rat tail collagen (Gibco, A1048301) was neutralized on ice in the presence of 10X DMEM with 0.1N NaOH such that the final concentration of the collagen is 1 mg/mL. Pellet of clusters was resuspended in 50 μL of neutralized collagen and seeded in 8-well chambered cover glass (Eppendorf 0030742036) and supplemented with defined medium. 3D cultures were grown in a 37°C humidified incubator with 5% carbon dioxide.

### Processing of 3D invasive clusters

3D invasion cultures were washed with phosphate buffered saline pH 7.4 (PBS) once after removing medium and fixed with 4% neutral buffered formaldehyde (Merck, 1.94989.0521) for 30 minutes. 2% glycine in PBS was used to neutralize traces of formaldehyde and blocked for one hour at room temperature with 5% BSA (HiMedia, MB083) in PBS+ 0.1% Triton X100 (HiMedia, MB031). After blocking, clusters were stained with 4’,6-diamidino-2-phenylindole (DAPI) (Invitrogen, D1306) and Alexa Flour 633-conjugated Phalloidin (Invitrogen, A22284) overnight at 4°C. Next day, cultures were washed with PBS+0.1% Triton X 100 for 10 minutes each three times.

### Laser scanning confocal microscopy and time lapse imaging

Processed clusters were imaged using laser scanning confocal microscope (Zeiss LSM 880) with system-optimized Z intervals. Images were analysed using Zen Lite software. Brightfield time lapse imaging was done using Olympus IX81 equipped with stage top incubator and 5% carbon dioxide (see Video 1). Imaging was done at 10-minute interval over 24 hours. Images acquired were analysed using ImageJ software (Schindelin et al., 2012).

### Computational model

#### Modelling environment

In order to simulate a biological milieu wherein cellular growth, proliferation, invasion and morphologies, consequences of multiple underlying processes of distinct length-scales and time-scales can be studied, a modelling environment is required which combines all those processes. The software package CompuCell3D (Swat et al., 2012) fulfils this purpose. Compucell3D is based on the Cellular Potts model, also known as Glazier-Graner-Hogeweg model that was designed to model collective behaviour of active matter (Chen et al., 2007; Sanyal and Glazier, 2006). This is done by calculating an energy function called Hamiltonian at each step of the simulation. Each cell is represented by the set of all Euclidian lattice sites or pixels sharing same cell ID. A rectangular Euclidian lattice has been used in all our simulations. The evolution of the model happens at each Monte Carlo step (MCS) which consists of index-copy attempts of each pixel in the cell lattice. Output of each MCS depends on the Hamiltonian calculation denoted by H (Figure 1B). The Hamiltonian in our model has two contributors which are affected by different properties of the cells and chemicals. The first contributor is the sum over all neighbouring pairs of lattice sites i and j with associated contact energies (J) between the pair of cells indexed at those i and j. In this term, i, j denotes index of pixel, s denotes cell index or ID, and t denotes cell-type. The δ function in this term will ensure that only the i≠j terms are calculated and also contact energies are symmetric. The contact energy between two cells can be considered as a term, which is inversely proportional to adhesion between those two cells. The second contributor is a function of the volume constraint on the cell, where for cell σ, lvol(σ) denotes the *inverse compressibility* of the cell, v(σ) is the number of pixels in the cell (*volume*), and *V*_t_(σ) is the cell’s *target volume*. For each cell, this term is governed by its growth equation. If any change in the Hamiltonian is negative at a given MCS for any configuration with respect to its previous one [Δ*H*□ = □(*H*_f_□−□*H*_i_)□<□0] then index-copy attempts of pixels resulting in that configuration will be successful. Otherwise, the attempt will be accepted with probability p = exp(Δ*H*□/T_m_). A default dynamical algorithm known as modified Metropolis dynamics with Boltzmann acceptance function is used at each MCS to move the system towards a low-energy configuration as MCS increases. The term T_m_ can be considered as temperature or magnitude of effective membrane fluctuations. In the model, the membrane fluctuation is kept high for cancer cells compared with matrix elements in order to strike a distinction between living and dead elements. Random movements of pixels leading to different transition probabilities at each MCS mimics the stochasticity present in biological systems.

**Figure 1:**
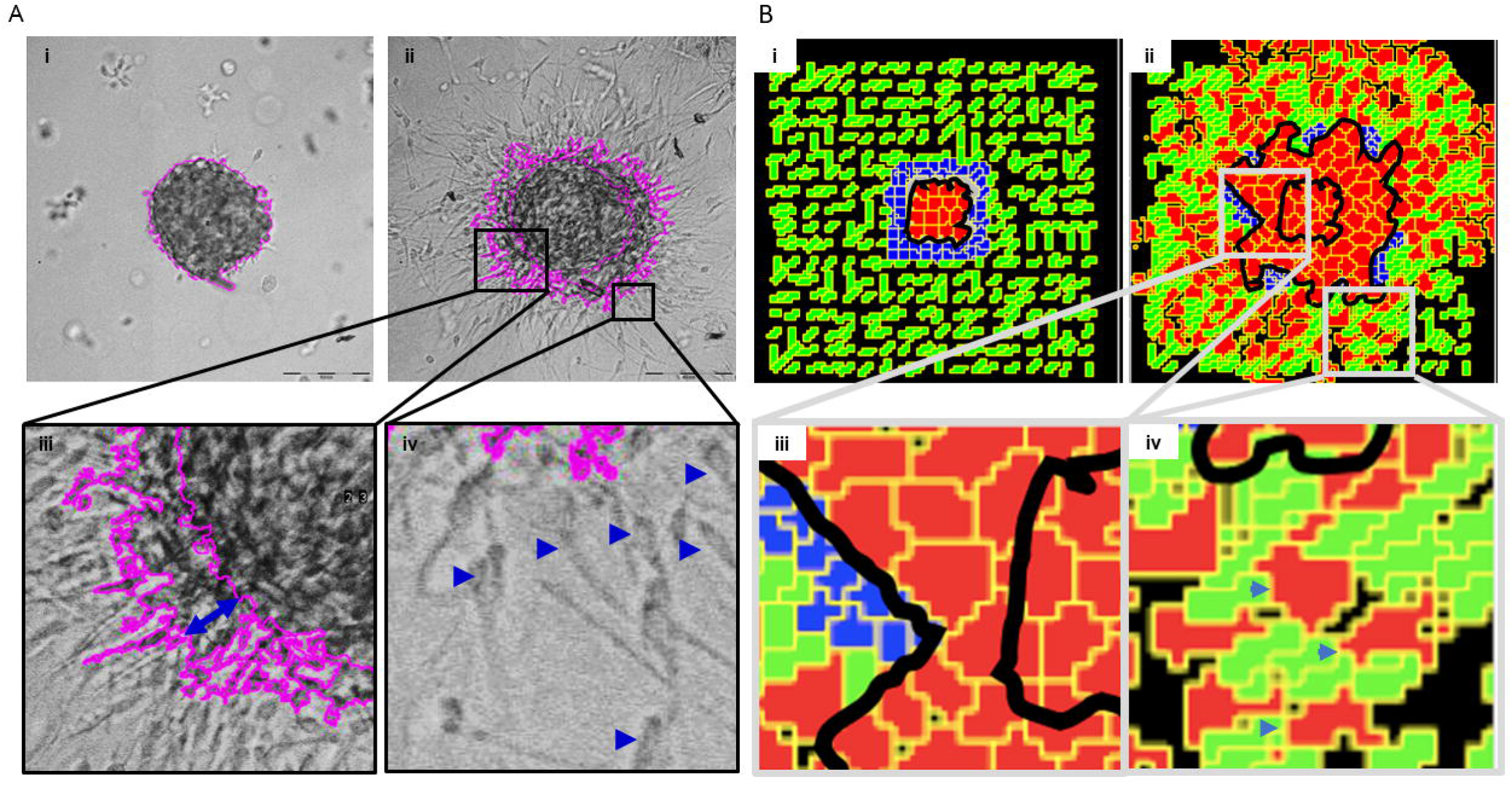
Schematic depiction of experimental system and computational model. (A) Early stages of breast cancer progression: Top left denotes the normal glandular (ductal/luminal) architecture of human breast. Top right denotes the pattern of breast epithelia undergoing ductal carcinoma-in-situ where normal epithelia are malignantly transformed, lose polarity and proliferate resulting in filling up of ductal lumen. Bottom right shows the architecture of invasive ductal/luminal carcinoma where transformed cells breach the basement membrane and invade into collagen-rich breast stroma. Bottom right shows how invasive breast cancer cells having traversed through stroma intravasate into blood/lymph vessels and metastasize to secondary organs. (B) Schematic workflow of 3D invasion assay used throughout the paper: Cells are first cultured on top of non-adhesive substrata in medium containing 4% rBM. Once cells assembled into clusters, the latter is embedded within Type 1 collagen scaffolds and cultured in serum-free conditions. (C) Governing equations used for setting up the computational model. The first equation calculates H which is Hamiltonian of the simulation: H determines the probability P associated with index-copy attempts, movements by generalized cells to minimize local effective energy through the default dynamical algorithm known as modified Metropolis dynamics. The equation for calculating growth rate shows it to be dependent on GH, Growth Factor concentration and g, which denotes nutrient availability. ECM remodeling follows the kinetics of reaction-diffusion (please refer to the appropriate sections in Material and Methods for a more detailed description of model construction). (D) Quantification of invasiveness of cancer cells in the computational model is performed using the minimal enclosing circle algorithm developed using MATLAB.

#### Model components

Cell & matrix orientation: Using a 2-dimensional computational model, several aspects of cancer invasiveness and tumor-associated 3-dimensional phenomena have been studied where the property of spherical symmetry of tumor morphologies was used to obtain the minimalistic setup(Jiao and Torquato, 2011). Our 2D simulations mimic experiments in which biological cells may require 3D space to allow certain interactions but in the computational model, only the properties associated with cells will play a role in determining the output irrespective of 2D or 3D. 2D simulations are computationally more efficient as it carries out exponentially less number of calculations for the whole system. Our model space is 100 * 100 *1 pixel in size where a group of cancer cells is initially located at the centre grid surrounded by ECM. Any element of the model that is required to actively participate through MCS pixel-copy attempts must be assigned a cell-type, as instructed, the laminin (‘BM’) and Type 1 Collagen (‘C1’) are assigned different cell-types along with cancer cells (‘CELL’). In the setup, clusters of cancer cells are surrounded by blob-like 2-layer cells of BM. The BM, in turn is surrounded by fibrillar collagen. To mimic in-vivo ECM architecture, BM is modelled as dense adhesive blob-like ‘cells’ similar to the lamina densa of basal lamina, whereas C1 is modelled as the interconnected fibres similar to Type 1 collagen. All components of the system have depth of 1 pixel in z direction so there is no overlapping of objects. A cell cannot cross another cell if it does not degrade it and without degradation the cell will be trapped in a zone due to steric hindrance by its surrounding environment or find ways to squeeze through small spaces in its vicinity which become accessible by random movement of that cell. In an initial configuration, cancer cells start as a rectangular objects of 16 unit volume (4*4*1pixels) spanning 14*14*1 unit volume at the centre (x,y= 43:57) of the simulation grid without any inter-cellular space. Tightly packed BM cells of 9 unit volume (3*3*1pixels) is then created around the cancer mass (x,y=37:63) having 2 layers of laminin cells separating it from C1. C1 is formulated around the cancer and BM structure throughout the whole grid with initial configuration of 4 unit volume cells (2*2*1pixels) with 2 pixel gaps in between them. In order to make the C1 fibrillar, a plugin is applied on the cells which elongate them in axis, random with respect to each other at 0.8 unit volume increment at each MCS till 5. The length-scale of components of ECM (BM and C1) is kept relatively smaller than cancer cells (Das et al., 2017). The lattices with no assigned cell-type or in other words the gaps or the free spaces are assigned cell-type ‘medium’ as a default of the Compucell3D syntax.

#### Contact energies (Differential Adhesion)

Compucell3D requires setting interactions between all cell-types in terms of contact energies. Higher contact energy values between two cell-types signifies lower adhesion or higher mechanical hindrance between them. This is denoted by the term J in the Hamiltonian (H). the contact energies are set in our simulation by qualitatively considering interactions between pairs of components of the experimental setup in terms of adhesion or repulsion. After running simulations with a range of values of each contact energy, from all the resultant combinations, an appropriate set of contact energies are taken. The contact energies that were established for model simulations included cell-cell, cell-BM, cell-C1, cell-medium. Values of these contact energies can be found in Table 1. The cell-medium contact energy can be thought in terms of cell-cell adhesion of cancer with proportional correlation. Higher cancer cell-cell adhesion will give higher cell-medium contact energy value. Across simulations, contact energies were established qualitatively motivated by transcriptomic findings including in, but not limited to (Kenny et al., 2007). For example, to mimic the decreased expression of E-cadherin in highly invasive cells, cell-cell contact energy was increased and cell-medium contact energy reduced

#### MMP-TIMP reaction diffusion system

Auxiliary equations in Compucell 3D are used to model chemical fields. These fields store the concentration information of a certain chemical at every location in the simulation grid. Two chemical fields, A and I are created which are governed by partial differential equations (PDE) based on reaction-diffusion dynamics of an activator and its inhibitor. These fields are incorporated with GGH algorithm to allow interaction between other simulation components and the fields. The governing equations for these two fields are:

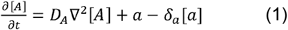

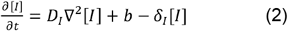

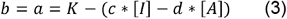

Where, [A], [I]: concentration values for fields A and I.

*D_A_, D_I_*: diffusion constants of A and I

*δ_A_, δ_I_*: degradation rates of A and I

a, b: secretion rate of A and I

t ≡ MCS

Default parameterizations, *D_A_*= 0.01, *D_I_* = 0.8, *δ_A_* = *δ_I_* = 0.003, K = 2.0, c = 4.0, d = 2.0

Here A is considered as the activated form of matrix metalloproteinases. Its activation (or secretion of the activated form i.e. ‘a’) is assumed to be dependent on its inhibitors (inversely) and on its own concentration (autocatalysis) in the form of equation-3. There are numerous variants of MMPs and TIMPs present in biological tissues (Brew and Nagase, 2010). Their production rates and inter-dependencies are still not known entirely for cancer cells so, a generalised form of MMP-TIMP interaction (A-I interaction of the model) is assumed in the light of reaction-diffusion dynamics (equation 1 & 2). The diffusion rate of MMP is set in the same range that is used by previous literature (Kumar et al., 2016) [that model has D= 1.0□×□10^−9^□cm^2^.s^−1^ = 0.025 pixel^2^ s^−1^, as 1mm=500pixels]. The difference in diffusion rates between the models (0.01 instead of 0.025) is due to different scaling of MCS with respect to time (s). All other parameters have been set based on previous literature and by optimisation of the model. The diffusion rate of I is set higher than A to generate localised ‘activator’ field and de-localised ‘inhibitor’ filed of reaction-diffusion system.

As all proteins have a lifetime, the degradation rate or decay constant associated with A and I in the model limits the spread of fields. (if there is no decay rate, initially secreted fields will spread the whole grid of the simulation as MCS increases without any further secretion). The Decay constant is assumed to be similar for A and I due to paucity of rigorous experimental analyses. In the model, A and I fields are secreted at the boundaries of all ‘CELL’ which come in contact with ECM. To initialise the A-I axis, random value of ‘a’ (range is from 0 to 4) is assigned to each cell which is in contact with ECM at MCS<5. CC3D package has forward Euler method-based PDE solver which was used to solve the PDE equations (Swat et al., 2012).

#### Matrix degradation and regeneration

Degradation of matrix is assumed to be dependent on the ratio of [A] over [I]. For each MCS, ECM cells access the concentration values of A and I fields at each of its pixels and depending on the ratio, those pixels are either degraded or remain unchanged. This degradation is threshold-based. Only pixels with the ratio [A]/[I] > 2.0, will be converted to ‘lysed’ form which is either C_Lysed (C1) or L_Lysed (BM) cell-type. The degraded matrix is assigned different cell-types by assuming different properties of degraded form of BM and C1 as the non-degraded BM and C1 also have different properties. As a cost of degradation, [A] and [I] are reduced at maximum 1.5 unit/MCS in pixels belonging to the ‘lysed’ cell-types. Matrix regeneration is incorporated into the model by conversion of C_Lysed and L_Lysed into C1 (given that cancer cells secrete predominantly fibrillar collagen-rich matrices) after 20 MCS from the degradation event associated with that ‘lysed’ pixel (Socovich and Naba, 2018; Yuzhalin et al., 2018). The regeneration of matrix is essential to eliminate unnecessary free spaces formed as an artefact of matrix degradation which takes the computational model closer to its experimental counterpart. All the ‘lysed’ cell-types are subjected to 0.1 volume decrease at each MCS to mimic dissipation of degraded matrix materials in-vivo.

#### Cellular growth and proliferation

Growth rate of ‘CELL’ is assumed be a linear combination of nutrient availability at cell boundaries and degradation of matrix. The growth equation is given by,

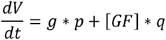

Where V = volume of ‘CELL’

g = measure of nutrient availability

[GF] = concentration of growth factor (GF) at centre of mass of ‘CELL’

p,q = constants

Two quantities, the common surface area of a cell with its neighbouring cells (k) and the total cell surface area (s) is accessed to calculate *g* in this equation as *g*=*(s-k)/40*. The denominator in calculation of *g* is due to 2-D nature of the simulation as a cell can be surrounded by other cells only in xy plane and not in z axis. The scaling of that extra cell surface area without any neighbouring cells in z axis is provided by the denominator. Another contributor of the growth function is *[GF]* which mimics the ECM-degradation dependence of growth and proliferation (Olivares et al., 2017). The ‘lysed’ cell-types are programmed to secrete GF at each of its pixel location where the diffusion constant is kept low (0.02) to localise this growth signal to areas of matrix degradation. p (=1/3) and q (=1/21) constant values are set according to the assumed weightage of the two variables in growth equation.

Cell division is incorporated into the cancer cells by a CC3D steppable called ‘MitosisSteppable’ with base function ‘MitosisSteppableBase’. If any ‘CELL’ reaches a threshold volume of 30 units then those cells will be divided in random orientation. The resultant two cells will have volumes half of its predecessor with all other properties kept same as the parent cell. In this model, growth rate is directly correlated to proliferation as it determines the volume of the cell to reach threshold for cell division.

#### Quantification: Invasion of morphology

The quantification for the spread or invasiveness of morphologies has been done in Matlab using minimal enclosing circle algorithm. Screenshots captured at different MCS from a simulation are used to track invasion of that model. A program was written where for a screenshot, the image is binarized with respect to ‘CELL’ which is represented by red colour. From that binarized image, centroids of all cells are accessed by the function ‘regionprops’. In order to find the smallest possible enclosing circle, two bits of information are required which are position of centre and radius of a circle which will encompass all the centroid positions. An arbitrary centre for the circle can be selected from which distances are measured to all the centroids. In the smallest possible enclosing circle, centre to centroid distance will be maximum for the furthest centroid that it needs to cover, and this distance will be the radius of the circle. The function ‘fminsearch’ is used with input of assumed centers and radii (maximum of center-to-centroid distance) which yields a center with minimized radius. Circle formed with this center and radius from ‘fminsearch’ will enclose all the centroids and will be the smallest possible circle to do so. All simulations have ‘CELL’ at the center of the grid as the initial configuration, so the centre of smallest enclosing circle can be assumed at the center of the that grid to start with, which is also the centre of screenshot; this optimises the programme. Running this program will yield smallest possible enclosing circle for screenshots at each specified MCS and the area of this circle is considered as the measure of invasiveness of that phenotype (Fig 1).

### Statistics

All the biological experiments were repeated three times independently. All the simulations were repeated at least 10 times and the data is represented as mean±S.E.M. Parametric students’ t-test was performed with Welch’s correction to estimate statistical significance.

## Results

### Breast cancer cells invade from rBM-like matrix to collagen-rich matrix concurrently across multiple spatial scales

In order to mimic the invasion of breast cancer cells *in vivo*, we designed a culture model, wherein MDA-MB-231 cells were allowed to form rBM-coated suspended clusters (See Materials and Methods; Fig S2 showing rBM is spatially limited to the surface of clusters). When such clusters were embedded in Type 1 collagen (Fig 2Ai) and the cultures were imaged in time lapse, the cancer cells rapidly migrated past the rBM barrier into collagen (Fig 2Aii). We observed spatially distinct but temporally concurrent modes of invasion, ranging from bulk motion where the cells moved centrifugally in an expansive and collective manner (Fig 2Aii bottom left inset), to mesenchymal migration of solitary cells with a slender cytoplasmic front and a nucleus-containing lagging end (Fig 2Aii bottom right inset). We here forward refer to this simultaneous deployment of distinct motility modes as multiscale invasion. Studies concerned with cancer cell migration investigate mechanisms underlying transitions between the modes; however the studies also note that these modes temporally coexist within histopathological sections of human tumors (Friedl and Alexander, 2011; Friedl et al., 2012; Krakhmal et al., 2015). Our experimental model successfully recapitulates the multimodal and multiscale 3D cancer cell invasion.

**Figure 2:**
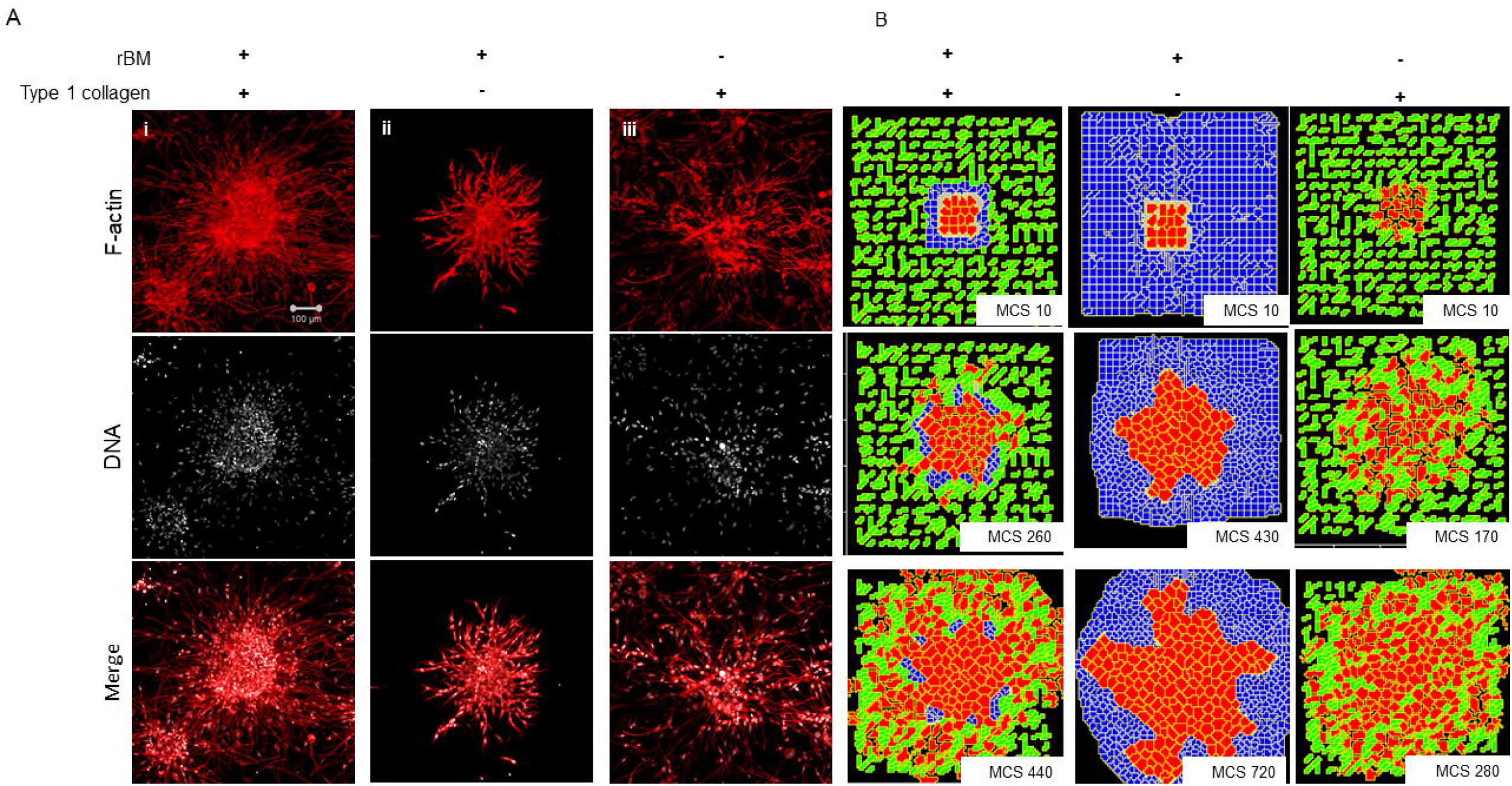
Multi-scale multicellular invasion of breast cancer cells in culture. (A) Representative phase contrast micrographs from time-lapse imaging of MDA-MB-231 cells showing multiscale invasion into fibrillar matrix. rBM-coated MDA-MB-231 clusters embedded within Type 1 Collagen (2Ai) invade into the latter within 24 h (2Aii). Cells show expansive migration (left inset; double-headed blue arrow shows the extent of collective migration between initial boundary (0 h) and final boundary (24 h) of the cluster (boundaries shown in pink). Single mesenchymal cells are also observed in Type 1 collagen (right inset; blue arrowheads). Scale bar: 200 μm. (B) Multiscale invasion exhibited by computational model. Initial pattern (2Bi; MCS10; cancer cells (red) packed within a basement membrane (BM)-like non-fibrillar matrix (blue) and further outwards by fibrillar collagen-like matrix (green; inter-fibrillar gap = 3 unit pixels)) and final pattern (2Bii; MCS 440) showing invasion of cells. In silico cells show expansive migration (left inset; bulk movement visible through the spatial gap between two black lines denoting boundary at initial MCS and final MCS). Single cells are also observed in non-fibrillar ECM.

We then sought to codify the minimal set of interactive cellular and ECM behaviors that could give rise such multiscale migratory behaviors of invading cancer cells. Using CC3D, we constructed a computational model, wherein for constrained set of values of cell-cell and -BM-like ECM adhesion, as well as upon invoking a reaction-diffusion (R-D) based remodelling kinetics of ECM, we observed multiscale invasion of cancer epithelia from a non-fibrillar to fibrillar in silico ECM microenvironment (Fig 2Bi represents the in silico cluster at Monte Carlo Step (MCS) = 10; Fig 2Bii represents the same cluster at MCS = 440; emergence of expansive collective invasion seen in Fig 2Bii bottom left inset, emergence of single cell invasion within the same culture seen in Fig 2Bii bottom right inset; see also appropriate sections in Materials and Methods for details of model construction). The use of an R-D-based modulation of ECM steady state was motivated by the morphology of the invasion phenotypes in our experimental assay, wherein invading cell populations were spatially separated by lateral zones of inhibition. In addition, the use of R-D based microenvironmental regulation and has strong precedence in the literature on cancer progression (Chaplain, 1995; Gatenby and Gawlinski, 1996; Roque et al., 2018; Zhang et al., 2018).

### Nature of the ‘stromal’ ECM may determine mode of cancer cell invasion

We sought to know whether the multiscale invasion of cancer cells was a function of the prototypical outwardly radial arrangement of cancer cells inside, a thin intervening layer of rBM and an outer presence of Type 1 collagen. To verify if the initial rBM coating was required for cluster shape and integrity, MDA-MB-231 cells were clustered in the absence of rBM. The cell clusters that formed had an irregular shape with ill-defined contours and were inherently unstable (Fig S3A and B showing irregular and regularly shaped clusters in the absence or presence of rBM coat, respectively). When rBM-coated MDA-MB-231 clusters were cultured entirely in rBM, clusters exhibited collective motility dynamics with most cells still attached to the kernel of the cluster (control multiscale invasion shown in Fig 3Ai; rBM-exclusive control shown in Fig 3Aii). Solitary invading cells were scarcely seen in the periphery. On the contrary, rBM-uncoated clusters upon embedding in Type 1 collagen gels, rapidly disintegrated into a small kernel and mostly single cells that exhibited mesenchymal single cell migration (Fig 3Aiii).

**Figure 3:**
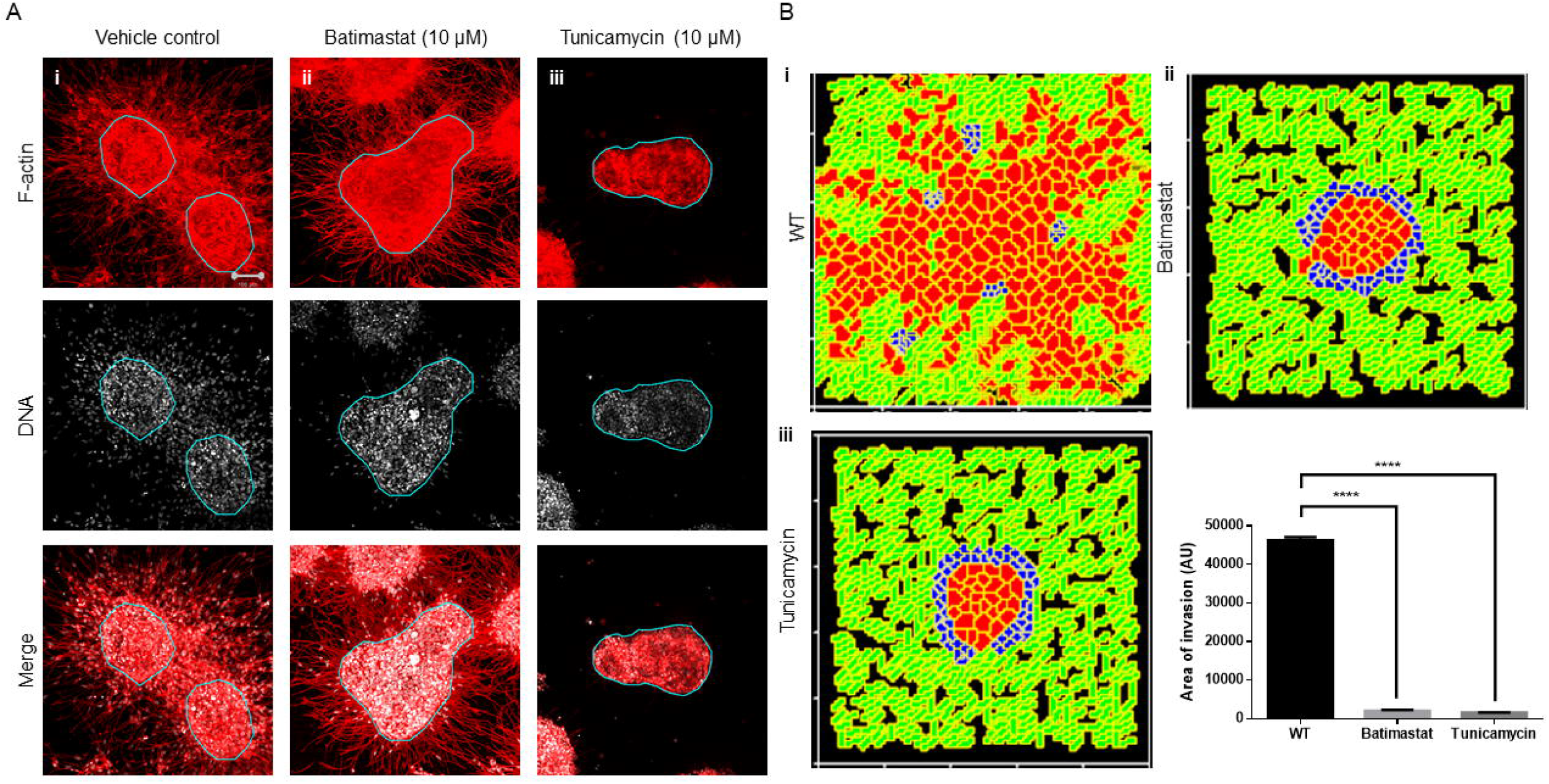
Single-matrix controls of models show simpler modes of invasion. (A) Maximum intensity projections of laser confocal micrographs of MDA-MB-231 cell clusters cultured within specific matrix milieu, fixed and stained for F-actin (using phalloidin; red; top row), DNA (using DAPI; white; middle row) and with both signals merged (bottom row). (i) rBM-coated clusters embedded in Type 1 collagen show multiscale invasion (left column) (ii) rBM-coated clusters embedded in rBM show collective or streaming migration of cells (iii) Uncoated MDA-MB-231 clusters in Type 1 Collagen show predominantly single cell invasion. Scale bar: 100 μm. (B) Representative images from simulations of invasion of cancer cells at early (top row) intermediate (middle row) and late MCS steps (bottom row). Simulations mimicking cells encapsulated within non-fibrillar and then fibrillar ECM exhibit multiscale invasion (left column). Simulations of cells cultured exclusively in non-fibrillar and fibrillar ECM show collective and single cell migration. Inter-fibrillar gap of C1= 3 unit pixels.

We used the phenotypic observations to further train our computational model and chose parametric combinations for i) contact energies of cell-cell, cell-rBM and cell-Type1 collagen interactions ii) reaction-diffusion-based remodelling of ECM and iii) proliferation and death of the cancer cells. We were able to successfully narrow down parametric combinations for which simulations mimicking ‘only rBM’- and ‘only collagen’ controls predicted predominantly collective and single cell migration respectively (Fig 3Bi represents control, Fig 3Bii shows emergence of collective invasion in an exclusive rBM-like non-fibrillar ECM environment, Fig 3Biii shows emergence of single cell invasion in an exclusive collagen-like fibrillar ECM environment. Since the parameter combinations were kept identical in the controls, the divergent phenotypes suggest that the identity of the stromal ECM and its spatial arrangement may determine the mode of outward migration of cancer epithelia.

### Metalloproteinase activity and N-linked glycosylation regulate multiscale invasion

We next sought to test our assumption that a locally auto-active regulation of ECM remodeling is essential for multiscale invasion. Matrix metalloproteinases (MMP) with their cognate lateral inhibitors: Tissue inhibitors of metalloproteinase (TIMP) are putative activator-inhibitor couples, given their diffusivity and nature of interactions. Treatment of cultures with a broad spectrum MMP inhibitor Batimastat resulted in an abrogation in transition of cells into the stroma, although the leading cytoplasmic head of cancer cells in the periphery of the cluster could still be visually discerned in the surrounding collagen (Fig 4Ai represents vehicle control; Fig 4Aii represents treatment with 10 μM Batimastat). This suggested that the transition of cancer cell nuclei across the rBM-collagen interface is dependent on protease-dependent remodelling of the stromal ECM. Interestingly, for amoeboid migration (which we have not investigated in our paper, see Discussion) nuclear softening has been proposed to be crucial for protease-independent migration (Das et al., 2019). Decreasing the activator levels within our computational model brought about a decrease in *in silico* migration of cells with sparse transitions into the fibrillar matrix environment (Fig 4Bi represents control; Fig 4Bii represents simulation that shows inhibition of invasion upon downregulating levels of activator A).

**Figure 4:**
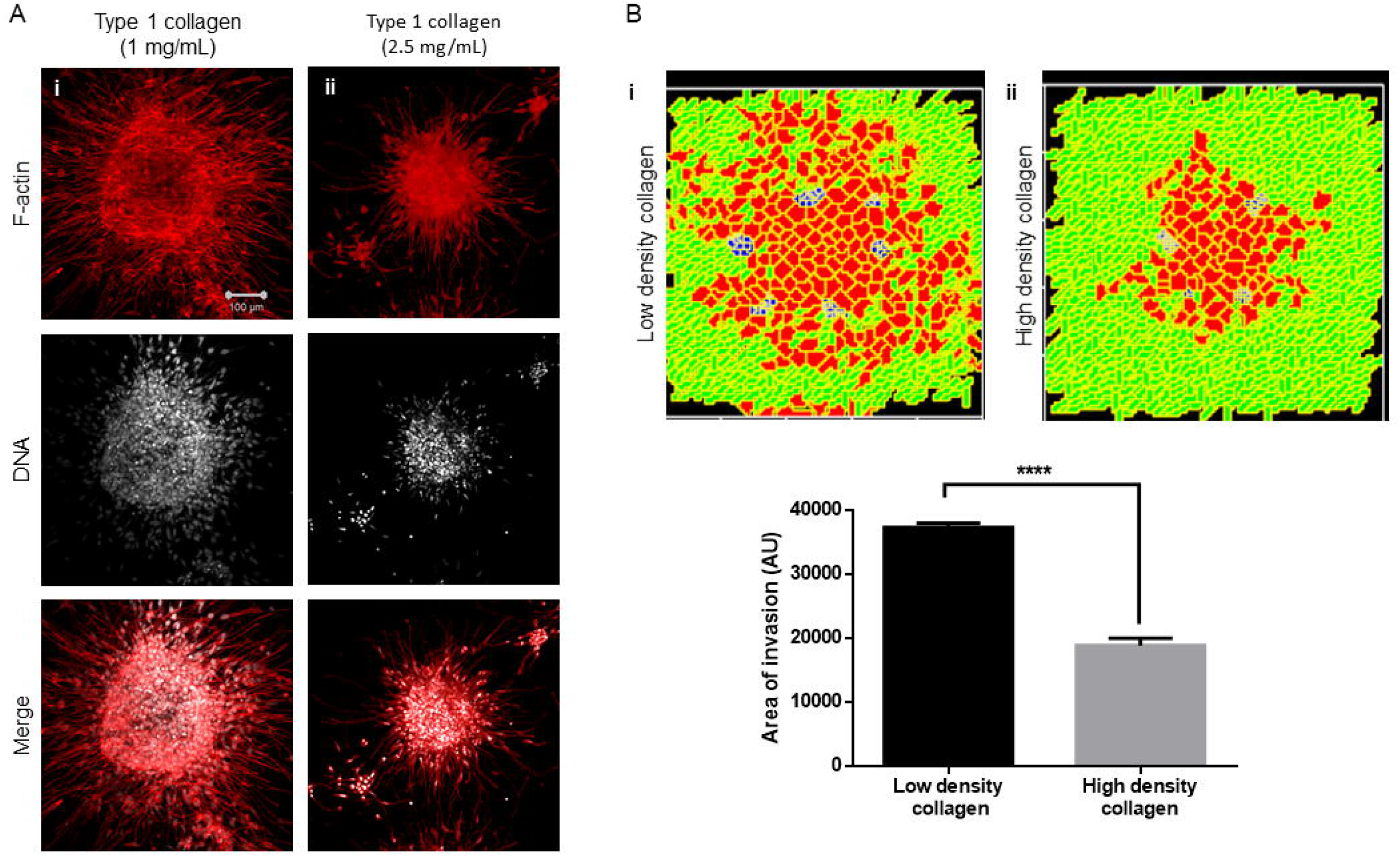
Inhibition of matrix metalloproteinase activity and N-linked glycosylation inhibits multiscale invasion. (A) Maximum intensity projections of laser confocal micrographs of MDA-MB-231 cell clusters cultured within specific matrix milieu, fixed and stained for F-actin (using phalloidin; red; top row), DNA (using DAPI; white; middle row) and with both signals merged (bottom row). (i) rBM-coated clusters embedded in Type 1 collagen treated with cehicle control DMSO show multiscale invasion (left column) (ii) Treatment with 10 μM Batimastat leads to inhibition of transition of cells to Type 1 collagen although cytoplasmic projections of cells in the periphery of the cluster are visible in the fibrillar matrix. (iii) Treatment with 10 μM Tunicamycin results in complete abrogation of multiscale invasion. (B) Simulations of control conditions (i), parametric variations analogous with inhibition of reaction-diffusion (ii) parametric variations analogous with inhibtion of cell-cell, cell-fibrillar ECM and reaction-diffusion (iii) at MCS590 Graph represents invasiveness of cells in simulations associated with 4Bi, ii and iii. Each bar represents mean +/− SEM **** denotes p-value <0.0001.

Second, we tested the role of cell-cell and cell-matrix interactions by treating our cultures with an inhibitor of N-linked glycosylation: tunicamycin. Tunicamycin affects the glycosylation and trafficking of cell surface proteins (Elbein, 1991). Molecules involved in cell adhesion such as cadherins and CAMs, are N-glycosylated. Moreover, while E-cadherin expression is epigenetically silenced in invasive MDA-MB-231 cells, N-cadherin is expressed and promotes their motility (Nieman et al., 1999). Treatment with tunicamycin does not alter the trafficking of N-cadherin but affects its function by interfering with its binding to catenin (Youn et al., 2006). Tunicamycin is also known to abrogate the matrix binding functions of integrins (Chammas et al., 1991). The effect of tunicamycin on metalloproteinases is context-dependent (Kim et al., 2010; Lee et al., 2019). Treatment with tunicamycin may increase the expression of MMPs, but due to associated ER stress and unfolded protein response, their secretion is inhibited (Duellman et al., 2015). Treatment of our complex experimental system with tunicamycin completely abrogated stromal transition of cancer epithelia (Fig 4Aiii). The cytoplasmic leading-edge extensions, likely mediated through outside-in integrin signalling, which were observed upon MMP inhibition, were also absent upon tunicamycin exposure.

In our computational model, the phenomenological equivalent of tunicamycin treatment would be to increase contact energies, and hence down-modulate adhesion between cells and matrices. Additionally, secretion of MMP and TIMP was also downregulated as part of the initial conditions for simulation. Upon doing so, we found impaired invasion of cells into the fibrillar in silico environment compared to control conditions (Fig 4Biii). We could also observe inhibition of invasion despite retaining the secretion of MMPs and TIMP but only under parametric combinations when the secretion rate of TIMPs exceeded that of MMPs (Fig S4). Our experimental and computational results suggest that adhesive interactions and local auto-active ECM remodelling dynamics operative within the invading milieu are necessary for stromal migration of cancer cells and inhibiting them significantly downregulates the latter (Fig 4Biv).

### Collagen density alters multiscale invasion

We next sought to test whether the arrangement of Type 1 collagen fibers surrounding rBM-coated clusters could regulate the nature of cancer cell migration. rBM coated clusters of MDA-MB-231 cells were embedded within a higher density of Type 1 collagen (2.5 mg/mL) scaffolds compared with control (1 mg/mL) (Fig 5Ai). The transition of cancer epithelia into high-density collagen was found to be attenuated (Fig 5Aii). Dense collagen may impede non-proteolytic migration of cancer cells allowing movement only upon mounting a protease-based degradation of ECM. In keeping with our experimental findings, in our computational model, we observe that all other parameters being kept constant, crowding the fibrillar ECM space with a higher density of collagen-like fibers decreased the migration of cells (Fig 5Bi represents control multiscale invasion; Fig 5Bii represents simulation in high density fibrillar ECM; Fig 5Biii shows statistically significant impairment of cellular invasion in the computational environment).

**Figure 5:**
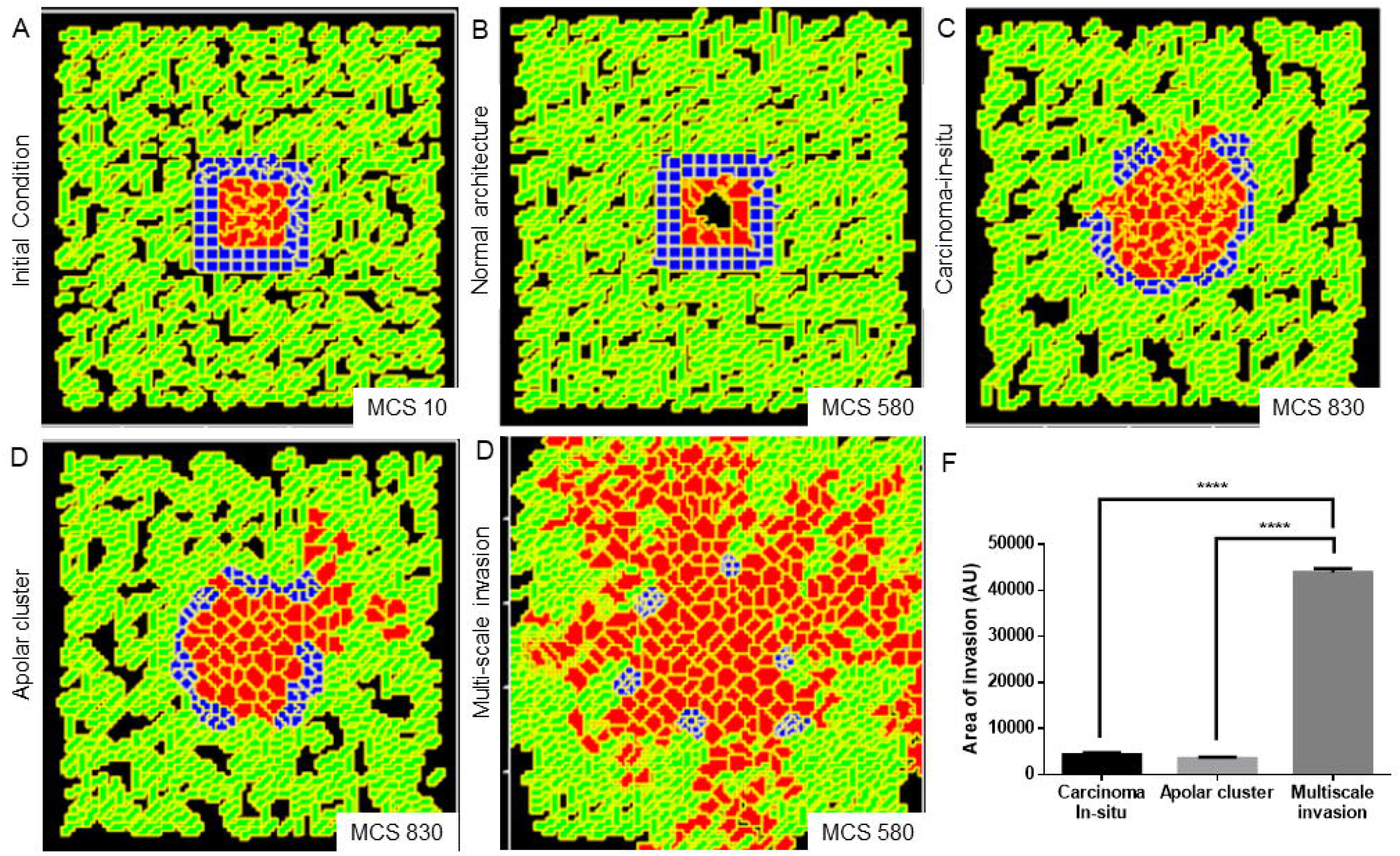
Increased collagen density impairs multiscale invasion. (A) Maximum intensity projections of laser confocal micrographs of MDA-MB-231 cell clusters cultured within specific matrix milieu, fixed and stained for F-actin (using phalloidin; red; top row), DNA (using DAPI; white; middle row) and with both signals merged (bottom row). (i) rBM-coated clusters embedded in 1 mg/ml Type 1 collagen show multiscale invasion (ii) rBM-coated clusters embedded in 2.5 mg/ml Type 1 collagen show impaired invasion of cells into surrounding high density Type 1 collagen (B) Simulations of control conditions (i), high density arrangement of fibrillar ECM showing impaired migration of cells at MCS 540. Graph represents invasiveness of cells in simulations associated with 5Bi and ii. Each bar represents mean +/− SEM ****denotes p-value <0.0001.

### Diversity in morphological phenotype can be explained by variation in interplay between cell adhesion and reaction-diffusion

Our computational model, trained on controls, successfully predicted the consequences on the phenotype of various perturbations. We asked whether it could also accommodate, with suitable changes in its formalism, the possibility of formation of homeostatic non-malignant phenotypes as well as precancerous and sub-invasive phenotypes? If so, what changes in the underlying coarse-grained physical mechanism could be responsible for those?

We obtained a non-invasive homeostatic lumen-containing phenotype (Fig 6A represents phenotype at MCS = 10; Fig 6B represents emergence of the phenotype at MCS = 580) by assigning the cells within our *in-silico* framework, certain properties similar to noncancerous ductal epithelial cells: basement membrane-regulated survival of the cells. By simply implementing the rules that 1. cells that are not anchored to the BM-like non-fibrillar ECM die (Frisch and Francis, 1994) and 2. cells anchored to the fibrillar ECM remain quiescent (Spencer et al., 2011), we were able to achieve growth-restricted lumen-containing acini-like structures, that resemble the structures formed by the non-malignant cell line HMLE in 3D (Fig S2A). In silico phenotypes similar to the precancerous carcinoma-in-situ like condition, which comprises filled multicellular masses of cells (similar to the mass morphology (Kenny et al., 2007)) (Figure S2B shows MCF7 cells forming similar architectures within our 3D assay) could be observed by increasing intercellular and cell-rBM adhesion (Fig 6C). A more sub-invasive morphology, which resembles the precancerous phenotype, but within which cells have lost their polarity and could give rise to indolently progressive tumors, has been referred to as ‘grape’ (Kenny et al., 2007). We simulated outcomes resembling this phenotype upon further relaxing the intercellular and cell-matrix adhesion (Fig 6D). It is crucial to note for simulating both the precancerous and indolent progression phenotypes, the reaction-diffusion-based ECM remodelling network was not deployed. Invoking the same and decreasing intercellular and cell-rBM adhesion brought about multiscale invasion in simulation (Fig 6E). Comparison of invasiveness between the simulations of three cancerous morphologies (Fig 6F) reveals that multiscale migration exhibits the highest invasiveness followed by the indolently growing cluster phenotype and in turn by the precancerous morphological phenotype.

**Figure 6:**
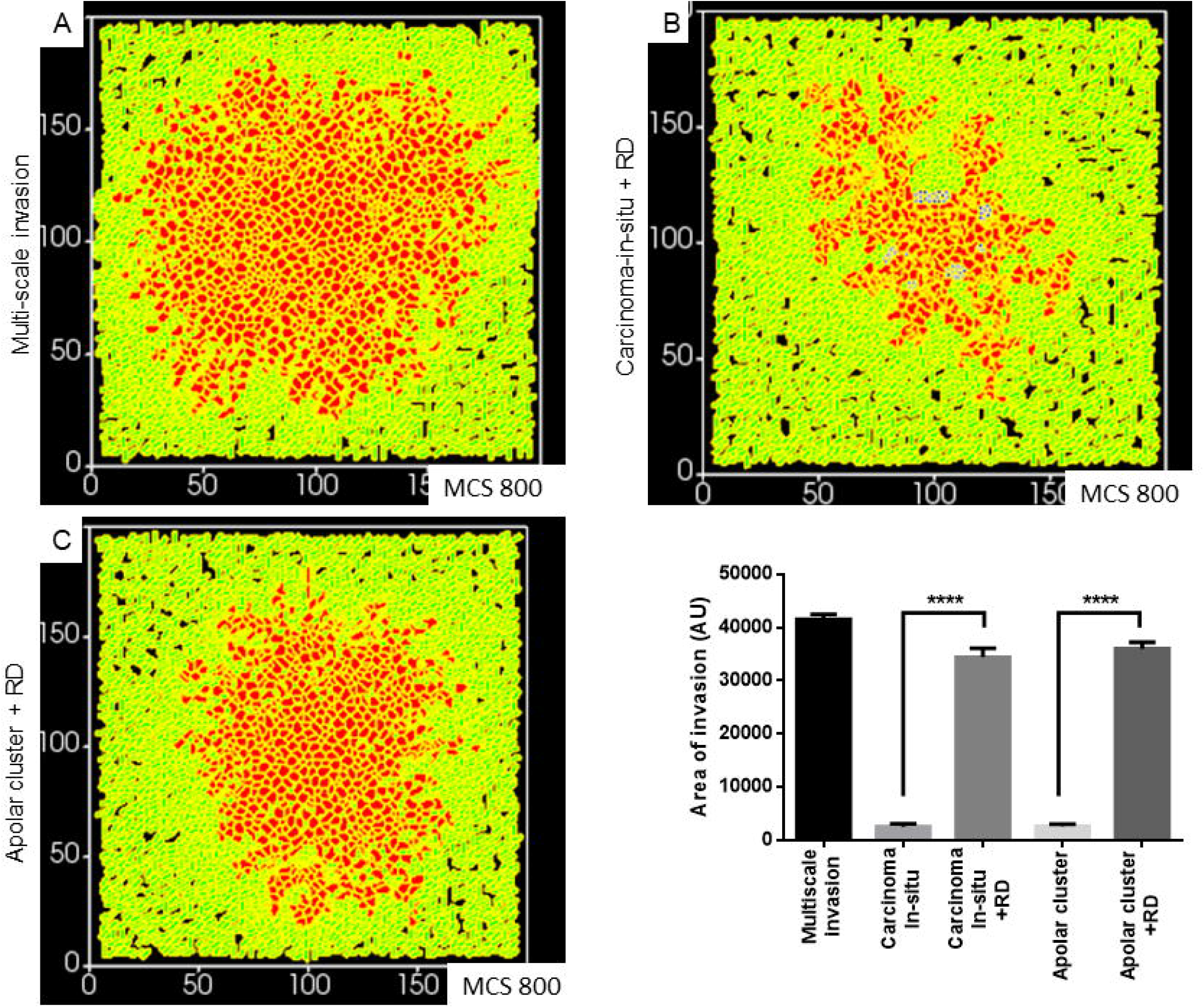
Parameter variation in the computational model can simulate homeostatic, precancerous and indolently cancerous phenotypes. (A) Initial configuration of all components of the computational model at MCS 0 (B) Simulation of a homeostatic growth-arrested phenotype with a central lumen obtained upon imposing a non-fibrillar ECM-based rules for regulation of cellular quiescence and death of anchored and detached cells respectively (C) Simulation of a carcinoma-in-situ-like phenotype obtained by maintaining high values of cell-cell and cell-non fibrillar ECM adhesion (D) Decreasing cell-cell and -ECM adhesion in simulation leads to a phenotype that shows further loss of polarity (as evidenced by a roughness in the outer contour of the clusters) and occasional sub-invasive single cell phenotypes (E) Further decreasing cell-cell and -matrix adhesion and deployment of a reaction-diffusion-based kinetics of ECM remodeling leads to multiscale invasion. (F) Quantification (bottom right) of the invasiveness of cells from simulations of homeostasis, carcinoma-in-situ, apolar clusters and multiscale invasion. Sacel bar: 100 μm. Each bar represents mean +/− SEM **** denotes p-value <0.0001.

Finally, we asked whether a decreasing gradient of cell-cell and cell-rBM adhesion was required for increased invasion as predicted by our simulations. Could merely deploying the reaction-diffusion-based ECM remodelling at higher adhesion regimes bring about greater invasion? Simulating diffusion-driven instability in ECM degradation in the context of the precancerous adhesion parameter values resulted in increased invasion that was exclusively collective and expansive (Fig 7A represents multiscale invasion; Fig 7B represents exclusively collective invasion upon simulating reaction-diffusion in the context of precancerous adhesion parameter values), and phenocopies the ónly rBM-like in silico morphology (see Fig 3Aii). On the other hand, simulating the same in the context of the adhesion regimes cognate to sub-invasive clustered morphologies did result in multiscale invasion (Fig 7C). It is to be noted however that the invasion seen in Fig 7B and C was significantly lesser than 7A but more than when in such same phenotypes and reaction-diffusion-based ECM modulation was off (Fig 7D) Our results implicate a threshold that lies between the precancerous and clustered adhesion regimes; the lower the cell- and matrix-adhesion, the greater the invasion.

**Figure 7:**
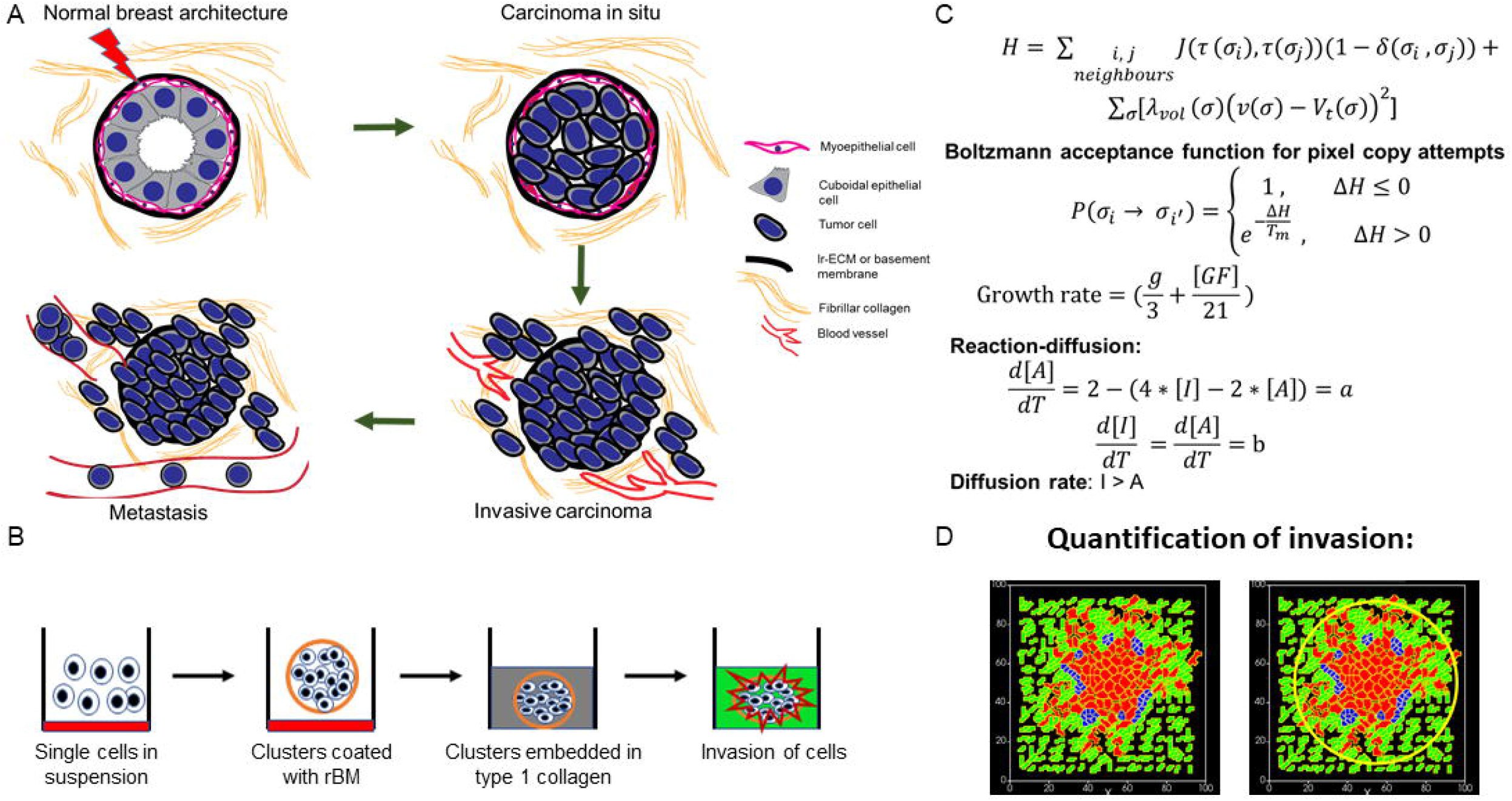
Simulations of the deployment of reaction-diffusion-based kinetics in carcinoma-in-situ and sub-invasive cluster phenotypes. (A) Simulations, within which regulation of the ECM was modeled using reaction-diffusion kinetics in parameter regimes of adhesion cognate to carcinoma-in-situ phenotypes, predict collective but not single-cell invasion (B) Simulations, within which regulation of the ECM was modeled using reaction-diffusion kinetics in parameter regimes of adhesion cognate to sub-invasive apolar cluster phenotypes predict multiscale invasion (C) Simulations for these results are done in 200*200*1 grid by keeping initial configuration similar to 100*100*1 grid. (F) Quantification of the invasiveness of cells from the above simulations in comparison with control multiscale invasion. Each bar represents mean +/− SEM **** denotes p-value <0.0001.

## Discussion

In this paper, we adopt a coarse-grained systems-theoretical approach towards the exploration of the mechanisms of stromal invasion of breast cancer epithelia. We designed an experimental organo-and patho-typic culture setup wherein not just the three-dimensional behaviour of cancer cells could be studied, but their transition from nonfibrillar (BM-like) to fibrillar (collagenous) ECM environments, as occurs in vivo could also be investigated. Using this assay, we observe this epithelial transition both as multicellular collectives and as single mesenchymal cells. In contrast, embedding cells in either (but not both) rBM and collagen (as controls) resulted in predominantly discrete collective and single cell migration respectively. Our observations imply that the complex multi-matrix nature of the assay presented here emulates in vivo invasive behavior to a better extent than existent single matrix assays.

Our experimental framework led to the construction, in parallel, of a computational model, whose parameters were trained on the phenotypic outcomes of various experimental controls. The design of the computational model takes inspiration from the concept of dynamical patterning modules (DPMs), autonomous heuristic agents that connote discrete physicochemical phenomena, such as adhesion, differential sorting, reaction-diffusion, polarity etc (Newman and Bhat, 2008, 2009). DPMs, when deployed singly or in combination, are useful for understanding the transformation of cellular patterns in distinct ways. DPMs has been used to investigate mechanisms of developmental morphogenesis in plants and animals (Benitez et al., 2018; Hernandez-Hernandez et al., 2012; Niklas and Newman, 2013). In addition, an DPM-based understanding of the evolution of development provides an explanation of how body plans of animals showed an accelerated period of origination (known as the Cambrian explosion) followed by a relative stasis (Newman et al., 2009).

In the specific context of breast cancer invasion, using DPMs, we have been able to treat much of the intracellular genetic repertoire and its associated dynamics (mutation and epigenetic regulations) as a black box. Instead, we concentrate on phenotypic traits that manifest at the spatial scales of cells and multicellular populations. We then asked whether specific combinations of parameters pertaining to these traits are permissive to the diversity of morphologies and cellular patterns seen in breast cancer progression. Given discrete assumptions that are confirmed by experiments, the same framework could give rise to phenotypes exhibited by non-malignant, malignant but non-invasive, sub-invasive and aggressively invasive malignant cells. In case of noncancerous cells, their quiescent and lumen-containing architecture was dependent on adhesion to basement membrane matrix; inability to do so resulted in anoikis (Bissell et al., 2002; Frisch and Francis, 1994; Furuta et al., 2018; Schwartz, 1997). The model predicts that the transition from homeostatic to a precancerous carcinoma-in-situ-like (DCIS) structures involves anchorage-independent survival and division. The transition from DCIS-like states to sub-invasive phenotypes that are characterized by complete loss of cell polarity involves a decrease in adhesion, both intercellular and between cells and BM-like matrices. On the other hand, the transition from sub-invasive phenotype to a full blown invasive multiscale phenotype is predicted to be achieved through specific interplay between decreased cell-cell and -matrix adhesion and reaction-diffusion-based cross-modulation between regulators of ECM remodelling, with neither physical process being sufficient by itself to bring about the phenotype. The computational model upon being asked to deploy reaction-diffusion in the presence of high cell-cell and -matrix adhesion predicted an exclusively collective invasion phenotype. The latter resembles morphologies obtained when cancer cell clusters are cultured in rBM scaffolds in the absence of Type 1 collagen. This suggests that the progression between two given morphologies can be achieved through distinct and dissimilar trajectories in parameter-space.

Reaction-diffusion-based mechanisms have been proposed to regulate the spatial patterning of iterative structures in development such as hairs, feathers, and digits (Glimm et al., 2014; Raspopovic et al., 2014; Sick et al., 2006). This occurs through interaction between an autocatalytic mediator of a morphogenetic step and its inhibitor (Gierer and Meinhardt, 1972; Meinhardt and Gierer, 2000). Both the mediator and its inhibitor are as per the R-D formalism, expected to be diffusible in nature (Turing, 1990). Their interaction would lead to spatial foci of morphogenesis separated by lateral zones of inhibition. It is reasonable to hypothesize the mediator to be a negative regulator of a morphological trait and its inhibitor to therefore antagonize the mediator’s inhibition of morphogenesis. Matrix metalloproteinases (MMP) and Tissue inhibitors of Matrix metalloproteinases (TIMP) are exemplars of such processes. They have been shown to play significant roles in mammary gland branch patterning (Wiseman and Werb, 2002). Their interaction dynamics in the context of mammary morphogenesis and elsewhere has been proposed to act through reaction-diffusion (Grant et al., 2004; Hoshino et al., 2012; Kumar et al., 2018; Skaalure et al., 2016).

A brief survey of expression patterns of genes across multiple cell lines grown on top of rBM matrices provides support for our predictions (Kenny et al., 2007). Cell lines exhibiting sub-invasive and invasive morphologies exhibit a progressive decrease in E-cadherin expression for which experimental support is available (Hiraguri et al., 1998). Cell lines with sub-invasive morphologies showed decreased levels of β1 integrin, which participates in multiple integrin heterodimers that bind to laminin. Invasive cells specifically expressed an aberrantly glycosylated levels of a β1 integrin (the consequences of glycosylation of β1 integrin have been reviewed in (Bellis, 2004)). Invasive cancer epithelia are known to express matrix metalloproteinases to a greater extent than untransformed cells: MDA-MB-231, for example, shows high levels of multiple MMPs as well as TIMP, relative to poorly invasive MCF7 cells (Bachmeier et al., 2001; Balduyck et al., 2000).

The modelling approach we have used successfully distinguishes between collective and single-cell growth dynamics However it is not able to parse mesenchymal versus amoeboid motilities. This is because we have modelled cells as bounded units that show little change in shape as they move. We aim to overcome this limitation in the future, by constructing multicompartment cells wherein intracellular cytoskeletal dynamics will be incorporated and will also be allowed to respond to inhomogeneities in ECM patterns. Our black-box approach also assumes a direct intracellular linkage between the various extracellular phenomena that mediate invasion. The introduction of signalling as a means of linking adhesion, proliferation, motility and ECM remodelling, and the (non)linear dynamics associated with the links would further enrich our understanding of the coordination between the diverse cellular phenomena in future efforts. In our computational model, cells proliferate copiously. On the other hand, our culture assays are grown for 24 to 36 h; cell proliferation can at best construed to play a mild role in the overall invasion. These two observations are not inconsistent with each other though; proliferation is also observed in cultures grown for longer time periods but does not alter the pattern of invasion that has been initially set by cell migration.

3D pathotypic cultures from patient cells/organoids are increasingly being considered as standards for personalized therapeutic strategies (Hagemann et al., 2017; Pauli et al., 2017). Their ability to prognose radio- and chemoresistance and match the results of patient derived xenograft models is backed up by a burgeoning body of literature (Gilles et al., 2018; Hubert et al., 2016; Zeeberg et al., 2016). Most of these culture setups lack a stromal compartment. The addition of the latter, as we have done in our assay, may prove to be a useful spatial milieu wherein the effect of immunotherapeutic interventions is tested. Our experimental breast cancer model can also be adapted for other cancers wherein the effect of stromal constituents on multiscale invasion of transformed epithelia may be studied and targeted.

## Acknowledgements

We thank Debayan Dasgupta for help with SEM. We also thank Divisional bioimaging facility, Biological science division at IISc. DP (Pally) acknowledges IISc for fellowship. DP (Pramanik) acknowledges KVPY for scholarship. RB acknowledges funding from CSIR (0412) and SERB DST Early Career Grant (1586) and the IISc-DBT partnership.

