## Supplementary figures and images for "An interplay between reaction-diffusion and cell-matrix adhesion regulates multiscale invasion in early breast carcinomatosis"

### Supplementary File 1

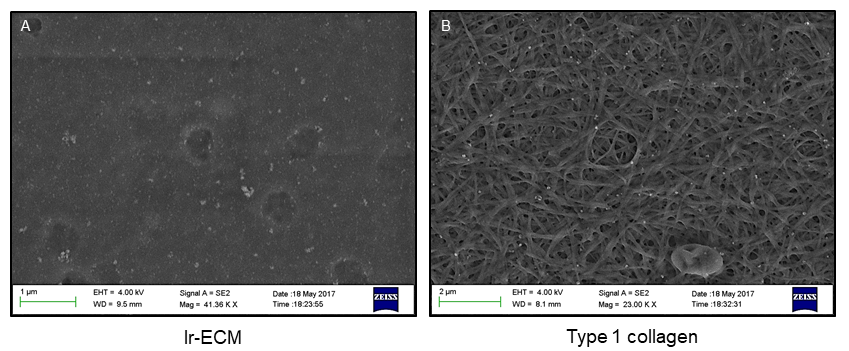

### Supplementary File 2

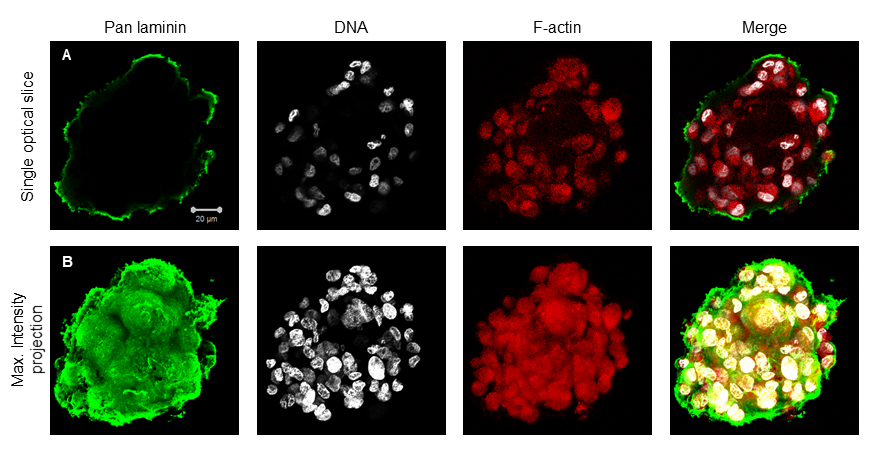

### Supplementary File 3

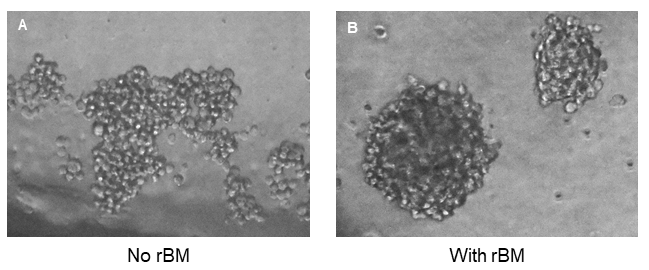

### Supplementary File 4

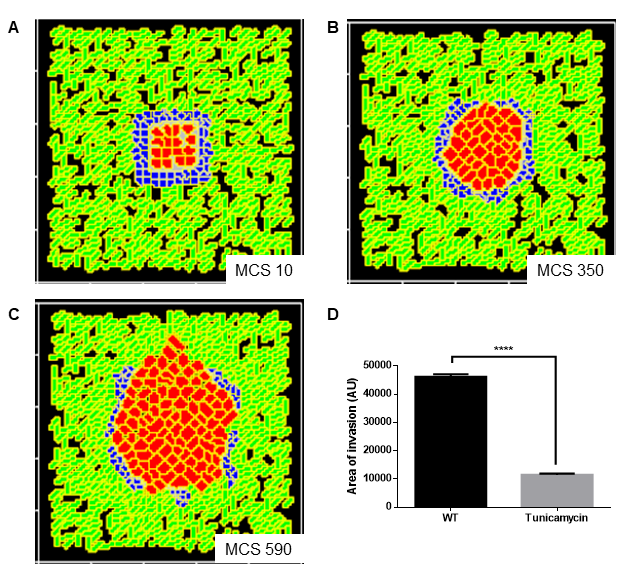

### Supplementary File 5

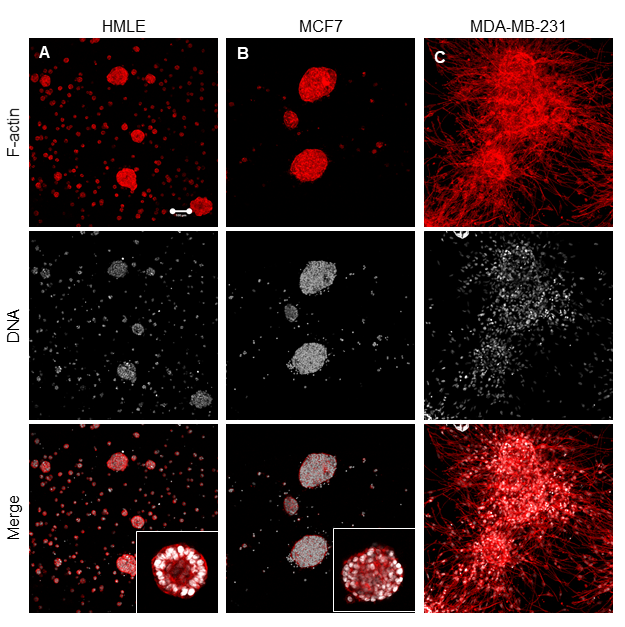
