## Supplementary Legends for "An interplay between reaction-diffusion and cell-matrix adhesion regulates multiscale invasion in early breast carcinomatosis"

Video 1: **Time lapse imaging of invasive cancer cell clusters in type 1 collagen.**

rBM coated MDA-MB-231 cell clusters embedded in type 1 collagen showing multiscale invasion. Scale bar: 200 μm

Figure S1: **Scanning electron micrographs of ECMs used in assays.**

(A) SEM micrograph of rBM shows sheet-like or non-fibrillar architecture. (B) SEM micrograph of polymerized Type 1 collagen shows fibrillar architecture.

Figure S2: **Presence of rBM on the surface of the MDA-MB-231 cell clusters**

(Top panel) Single optical slice of laser confocal micrograph showing the presence of rBM (stained with pan laminin antibody) exclusively on the surface of the clusters. (Bottom panel) Maximum intensity projection of the same showing surface staining of pan laminin on MDA-MB-231 cell clusters. (Green: pan-laminin; White: DNA; Red: F-actin) Scale bar = 20 μm.

Figure S3: **Phase contrast micrographs of MDA-MB-231 cells clusters cultured on non-adherent substrates.**

(A) Formation of rough edged and loosely formed clusters in the absence of rBM after 48 h of culture in suspension (B) Smooth boundary and tightly packed clusters are formed when the cells were cultured in 4% rBM containing medium for 48 h in suspension. Objective:10X

Figure S4: **Simulations of the deployment of reaction-diffusion-based kinetics in carcinoma-in-situ and sub-invasive cluster phenotypes**

(A) Simulations, within which regulation of the ECM was modeled using reaction-diffusion kinetics in parameter regimes of adhesion cognate to carcinoma-in-situ phenotypes, predict collective but not single-cell invasion (B) Simulations, within which regulation of the ECM was modeled using reaction-diffusion kinetics in parameter regimes of adhesion cognate to sub-invasive apolar cluster phenotypes predict multiscale invasion (C) Simulations for these results are done in 200*200*1 grid by keeping initial configuration similar to 100*100*1 grid. (F) Quantification of the invasiveness of cells from the above simulations in comparison with control multiscale invasion. Each bar represents mean +/- SEM **** denotes p-value <0.0001.

Figure S5: **Growth arrested, precancerous and multiscale invasive phenotypes**

Laser confocal micrographs of maximum intensity projected images of rBM-coated cell clusters embedded in Type 1 collagen. (A) Immortalized mammary epithelial cells (HMLE), form growth-arrested acinar-like structures. Merge inset shows a single acinar-like structure with a lumen (B) Non-invasive breast cancer cells (MCF7) show a carcinoma-in-situ-like phenotype. Merge inset shows a cell-filled MCF7 cluster. (C) Invasive breast cancer cells (MDA-MB-231) show multiscale invasion. Scale bar = 100 μm.
